# Endothelial cell Pannexin1 overexpression impairs ischemic stroke outcome in a sex-dependent manner

**DOI:** 10.1101/2025.02.07.636909

**Authors:** Maria Tomàs-Gracia, Amanda K Mauro, Colleen K Duffy, Eric Y Dai, Guleer Shahab, Christopher B Medina, Kodi S Ravichandran, Brant E Isakson, Miranda E Good

## Abstract

Ischemic stroke is a leading cause of morbidity and mortality. We have previously shown that deletion of endothelial cell (EC) Panx1 reduces ischemic stroke infarct volume and reduces cerebral arterial myogenic reactivity, which regulates cerebral blood flow. We hypothesized that EC Panx1 content dictates ischemic stroke outcome and thus increased EC Panx1 expression will worsen ischemic stroke outcomes due to exacerbated myogenic tone development and impaired cerebral blood flow recovery. To test this, we generated the Cdh5-Cre^ERT2+^ ROSA26-hPanx1^Tg^ mouse model that conditionally overexpresses the human isoform of Panx1 specifically in EC. We have found that cerebral myogenic reactivity is significantly increased with overexpression of EC Panx1 only in female mice, without alterations in peripheral vascular reactivity or blood pressure regulation. Similarly, we found that infarct size was increased and recovery of cerebral blood flow was reduced in female but not male EC Panx1 overexpressing mice. Our findings indicate a role for EC Panx1 as a mediator of ischemic stroke recovery. Furthermore, these data suggest a potential sex-dependent effect for EC Panx1, where females are more sensitive to increased EC Panx1 in cerebral vascular function and may provide a potential therapeutic target for the treatment of ischemic stroke in women.

## Introduction

In the US, around 800,000 people experience an acute ischemic stroke per year, costing an estimated $36 billion^1^. Acute treatment is still limited to thrombolytic therapy, with agents such as tissue plasminogen activator (tPA) or endovascular thrombectomy^2^. tPA, the only FDA-approved drug for acute treatment, aims to break down the occlusion in the cerebral blood vessel, thus preventing further damage, yet it neglects to reduce the extent of the injury post-reperfusion. In addition, its short window of effectiveness of about 3-4.5 hours after stroke onset and the associated hemorrhagic risks considerably limits its therapeutic ability^3^. Following an ischemic stroke, proper reperfusion through the cerebral vasculature and recovery of cerebral blood flow (CBF) can determine the initial extent of injury to the brain^4, 5^. However, the mechanisms dictating reperfusion into the ischemic region are incomplete. Women have been found to have an increased lifetime risk of stroke and stroke-related mortality even though risk of ischemic stroke increases with age in both men and women^2^. Given that the global incidence of ischemic stroke is projected to increase over the next 10 years and an increasing number of patients will survive a stroke with significant morbidity^6^, it is necessary to further explore new therapeutic approaches while elucidating underlying physiological mechanisms of ischemic stroke injury and potential sex-specific differences in these mechanisms.

CFB is largely regulated through myogenic reactivity, in which arteries constrict in response to increased luminal pressure^7, 8^. Since adequate restoration of CBF is crucial to recovery after acute ischemic stroke^4, 5, 9^, understanding the mechanisms of myogenic tone regulation and its impact in pathological states is key to reveal potential therapeutical targets. We and others have identified a role for purinergic signaling in regulating myogenic tone in the cerebral vasculature and ischemic stroke outcomes^10, 11^. P2Y4 and P2Y6 receptors have been shown to be important for development of myogenic tone in cerebral arterioles^10^. Additionally, we recently found that Panx1 channels were necessary for development of cerebral artery myogenic tone development^11^. Pannexin1 (Panx1) channels are ubiquitously expressed plasma membrane spanning proteins that form large conductance channels acting primarily as the purine-release channel^12^. Panx1’s regulated release of purines into the extracellular space activates downstream purinergic receptors, such as P2Y6 or adenosine receptors, thus playing a vital role in regulating purinergic signaling mechanisms^13, 14^.

Panx1, the primary purine nucleotide release channel, is expressed in most cell types, including EC, vascular smooth muscle cells (SMC), neurons, and glial cells^11, 15–17^. However, until our recent study^11^, no one had examined the functional role of Panx1 in the cerebral vasculature or the role of vascular Panx1 expression in ischemic stroke outcome. Our previous work identified endothelial cell (EC) Panx1 as a regulator of cerebral vascular function, where deletion of EC Panx1 reduced cerebral artery myogenic tone development but had no effect on peripheral mesenteric artery myogenic tone or systemic blood pressure^11^. Additionally, we found that the deletion of EC Panx1 significantly reduced post-ischemic stroke infarct volume^11^. These data suggested that EC Panx1 expression is necessary for cerebral myogenic tone and development of a post-ischemic stroke infarct; however, it is unknown if increased EC Panx1 exacerbates the detrimental effects of Panx1 in the cerebral vasculature post-ischemic stroke.

Here, we explored the effect of overexpression of human Panx1 in EC on myogenic tone, regulation of CBF and ischemic stroke outcomes in both female and male mice. We crossed our novel ROSA26-hPanx1^Tg^ mice^18^ with the Cdh5-Cre^ERT2+^ mice to inducibly overexpress the human isoform of Panx1 specifically in EC. Our data demonstrate that the overexpression of EC Panx1 increased myogenic tone development in cerebral arteries without altering peripheral vascular function only in females. Furthermore, increased EC Panx1 expression resulted in worse acute ischemic stroke infarct volume and impaired CBF recovery 24 hr post-reperfusion only in females. Together these data suggest that EC Panx1 content may dictates ischemic stroke outcome in females.

## Results

### EC Panx1 was overexpressed in cerebral EC in both male and female Cdh5-Cre^ERT2+^ Rosa26-Panx1Tg mice

Mice were made by crossing our Rosa26-human Panx1^Tg^ (R26-hPanx1^Tg^) mice^18^ with our Cdh5-Cre^ERT2+^ mice to create a mouse that conditionally expresses the human isoform of Panx1 with a FLAG tag (**Fig 1A**) specifically within EC. We confirmed the conditional expression of the flag tagged Panx1 within EC by staining posterior cerebral arteries (PCA) with an anti-Flag tag antibody in Cdh5-Cre^ERT2+^-R26-hPanx1^Tg^ (EC Panx1 OvExp) and Cre negative R26-hPanx1^Tg^ control (EC Panx1 Intact) mice (**Fig 1B**). To quantify the total expression level of Panx1 in EC within the brain, we isolated EC from the brain and performed western blots (**Fig 1C**). Panx1 expression was significantly increased in both female and male EC Panx1 OvExp mice with an ∼57% increase in female mice (p=0.0241) and ∼65% increase in male mice (p=0.0178) (**Fig 1D**).

**Figure 1.**
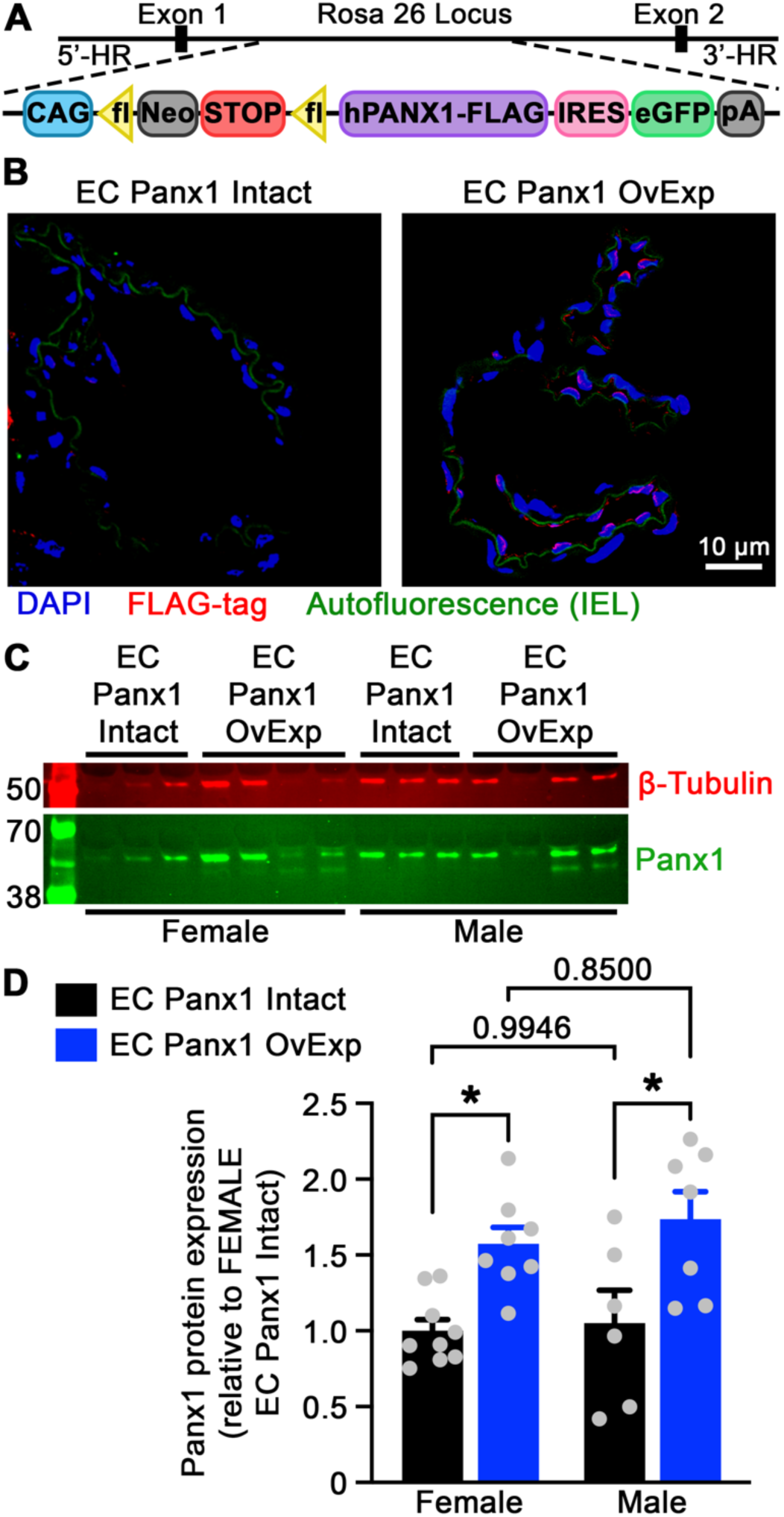
Panx1 is overexpressed successfully in ECs of cerebral arteries of both female and male mice. **(A)** Cartoon of the inserted human isoform of Panx1 (hPanx1) with a FLAG tag in the ROSA 26 locus to induce conditional overexpression of Panx1. **(B)** PCA from Cre^−^ R26-hPanx1^Tg^ control mice (EC Panx1 Intact) and Cre^ERTT2+^ R26-hPanx1^Tg^ (EC Panx1 OvExp) were stained for hPanx1 using the anti-Flag tag antibody (red). Internal elastic lamina is identified by its autofluorescence (green) and nuclei were identified using DAPI (blue). Western blot **(C)** and quantification **(D)** for Panx1 protein expression levels from ECs isolated from the brain of female and male Cre^−^ R26-hPanx1^Tg^ control mice (EC Panx1 Intact) and Cre^ERTT2+^ R26-hPanx1^Tg^ mice (EC Panx1 OvExp). Β-Tubulin (red) was used to normalize Panx1 expression (green) for quantification. Quantification of Panx1 protein expression is relative to levels in control (EC Panx1 Intact) female mice. EC Panx1 Intact: N = 9 females and 6 males. EC Panx1 OvExp: N = 8 females and 7 males. Two-way ANOVA with Tukey’s multiple comparisons test. *p<0.05.

### Overexpression of endothelial Panx1 did not alter <u>peripheral</u> myogenic tone, blood pressure or heart rate in female and male mice

To evaluate if the overexpression of EC Panx1 altered baseline peripheral and systemic vascular function we examined myogenic tone in mesenteric arteries and blood pressure in our EC Panx1 overexpressing and control mice. Similar to our results with deletion of EC Panx1^11^, we found no difference in myogenic tone development or passive diameter in mesenteric arteries between EC Panx1 overexpressing and control mice in both sexes (**Fig 2A-F**). Additionally, mean arterial pressure (MAP) and heart rate, obtained from radiotelemetry, were conserved across groups (**Fig 2G and H**).

**Figure 2.**
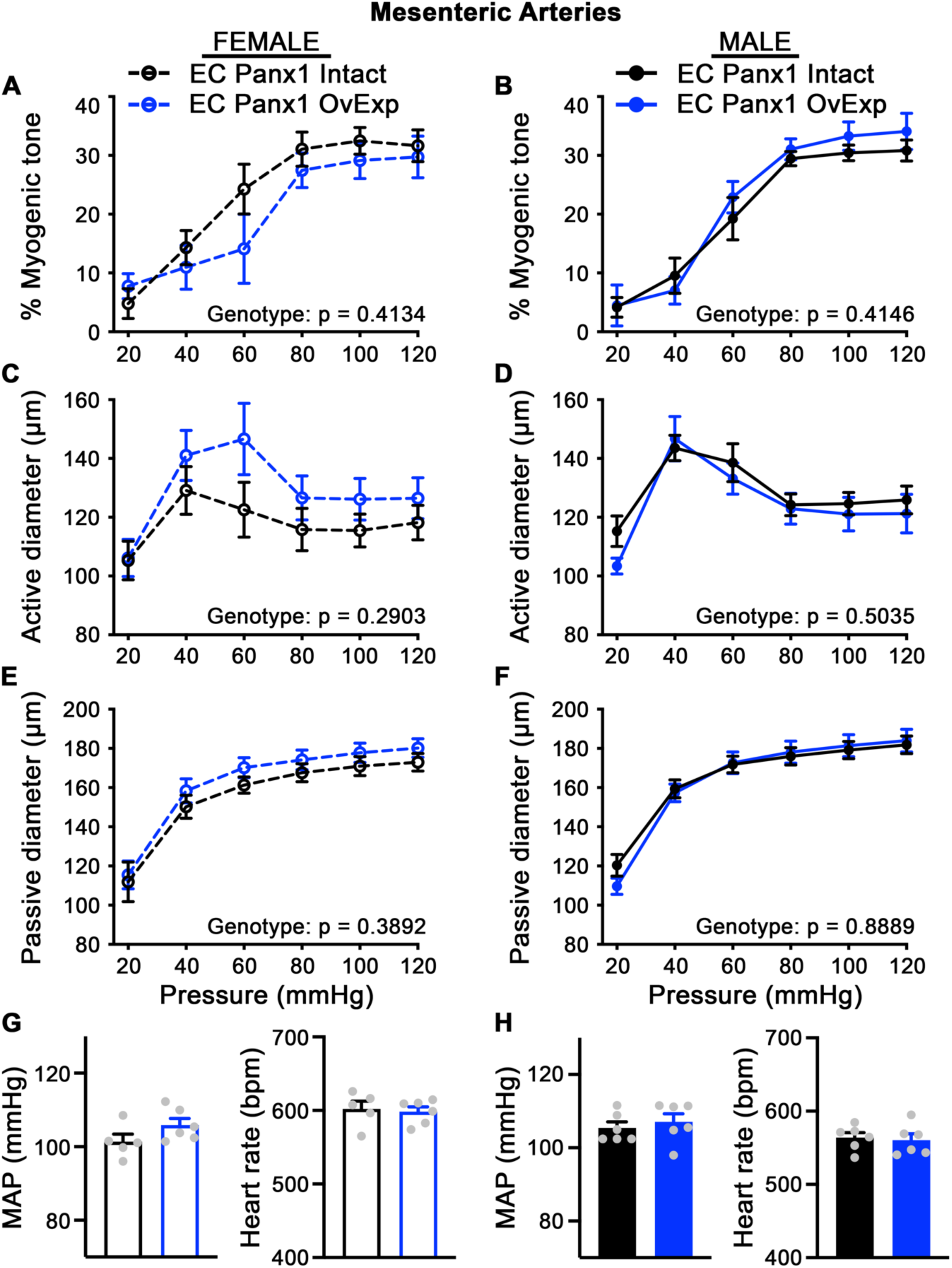
EC Panx1 levels do not impact peripheral vascular myogenic tone, systemic blood pressure and heart rate. Pressure myography was used to measure percent myogenic tone **(A-B)**, active diameter **(C-D)**, and passive diameter **(E-F)** of isolated third-order mesenteric arteries from female and male EC Panx1 overexpressing and control mice. EC Panx1 Intact: N = 6 females and 7 males. EC Panx1 OvExp: N = 6 females and 5 males. Two-way ANOVA with repeated measures and Sidak’s multiple comparisons post hoc test. Radiotelemetry was used to collect average mean arterial pressure (MAP) and heart rate from female **(G)** and male **(H)** EC Panx1 overexpressing and control mice. EC Panx1 Intact: N = 5 females and 6 males. EC Panx1 OvExp: N = 6 females and 6 males. Student’s T test. *p<0.05.

### Overexpression of endothelial Panx1 increased cerebral artery myogenic tone only in female mice

We next examined the development of myogenic tone in PCA and found that female EC Panx1 overexpressing mice had a significant reduction in their active diameter (**Fig 3C**) over the pressure curve with no change in passive diameter (**Fig 3E**), resulting in increased cerebral myogenic tone (**Fig 3A**) compared to control female mice. However, myogenic tone development of was not different in male EC Panx1 overexpressing mice compared to control mice (**Fig 3B**). The passive diameter was found to be significantly reduced in male EC Panx1 overexpressing mice (**Fig 3F**); although a mild, but not significant, decrease in active diameter (**Fig 3D**) resulted in no change in myogenic tone.

**Figure 3.**
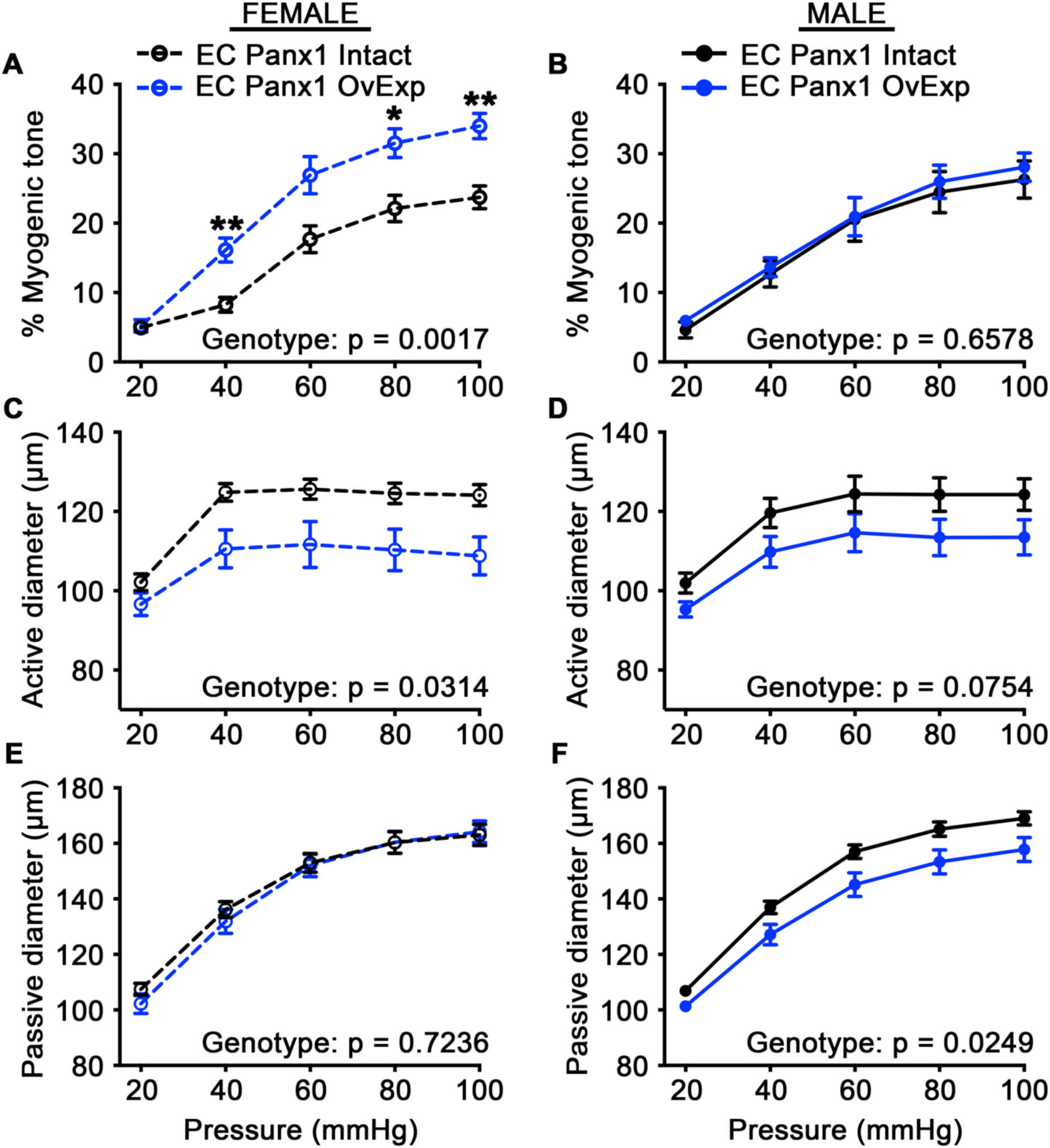
EC Panx1 overexpression increases myogenic tone in cerebral arteries only in female mice. Pressure myography was used to measure percent myogenic tone, active diameter, and passive diameter of isolated PCAs from female and male EC Panx1 overexpressing and control mice. EC Panx1 Intact: N = 9 females and 9 males. EC Panx1 OvExp: N = 11 females and 8 males. Two-way ANOVA/Sidak’s multiple comparisons post hoc test. *p<0.05; **p<0.01.

### Overexpression of endothelial Panx1 increased ischemic stroke infarct volume and impaired cerebral blood flow recovery only in female mice

To examine if increased myogenic tone altered ischemic stroke recovery in female EC Panx1 overexpressing mice, we performed a 45-minute unilateral middle cerebral occlusion (MCAO) followed by a 24 hr reperfusion. Baseline CBF was not different across genotypes and sexes (two-way ANOVA, sex: p=0.37 and genotype: p=0.21). To confirm successful occlusion of the MCA, cerebral blood flow (CBF) was measured at the end of the 45 min MCAO and compared to baseline CBF measured 24hr prior to MCAO. We found no difference in CBF reduction between genotypes or sex (two-way ANOVA, sex: p=0.54 and genotype: p=0.79) (**Fig 4A and C**), indicating a similar CBF reduction in ischemic injury. 24 hr after reperfusion, CBF was measured to assess the acute blood flow recovery in the ischemic hemisphere post-MCAO/reperfusion. Only female EC Panx1 overexpressing mice had a significantly reduced CBF recovery at 24 hr post-MCAO compared to their respective baseline CBF (p=0.0180) (**Fig 4G**). Male mice, both control and EC Panx1 overexpressing mice, recovered their CBF back to baseline levels (**Fig 4H**). Immediately after CBF imaging, brains were harvested and infarct volume quantified. In line with impaired CBF recovery, female EC Panx1 overexpressing mice had a significantly larger infarct volume compared to control female mice 24 hr post-reperfusion (p=0.0050) (**Fig 4B and E**). Male EC Panx1 overexpressing mice had a similar infarct volume compared to control male mice (**Fig 4D and F**) 24 hr post-reperfusion (p=0.34).

**Figure 4.**
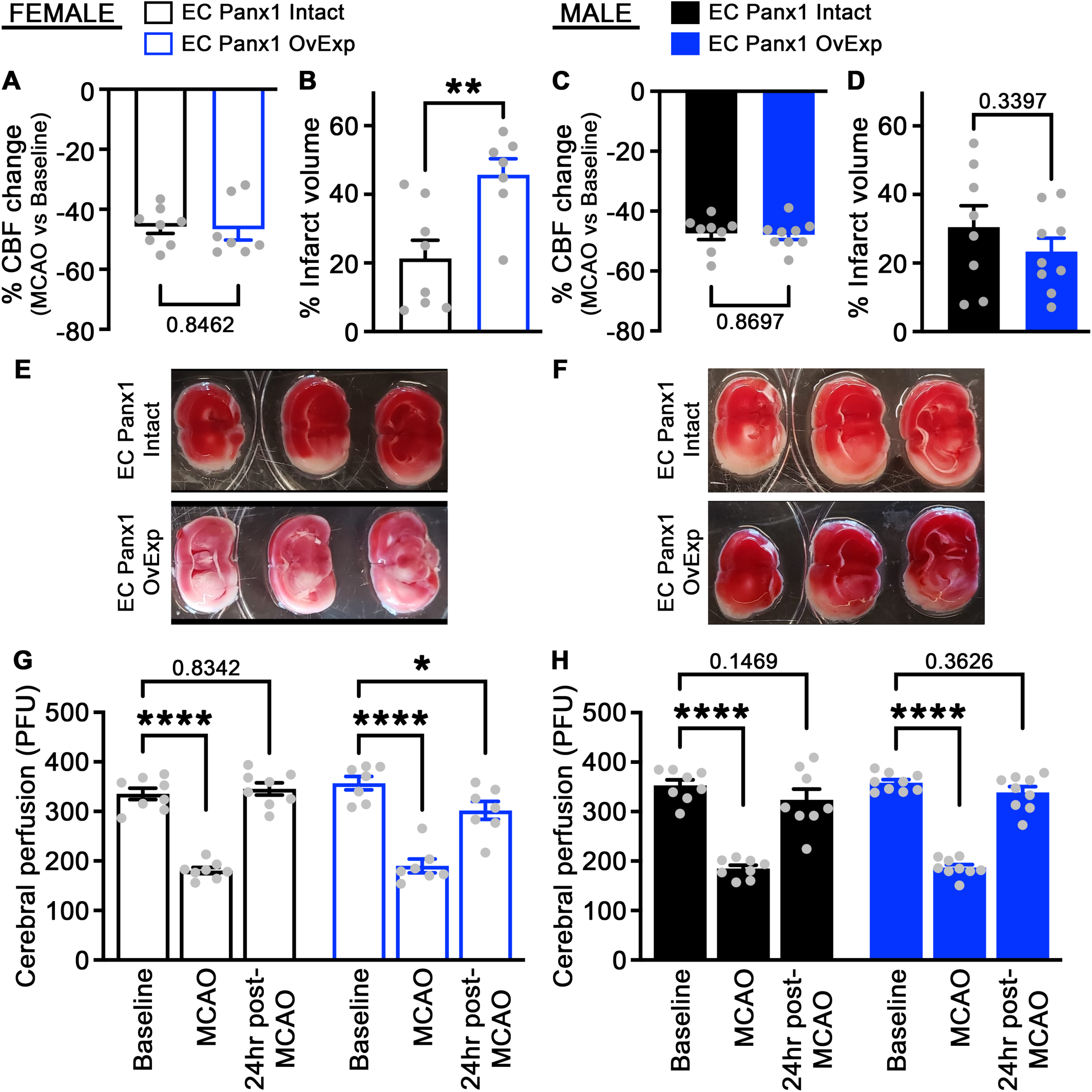
Overexpression of EC Panx1 significantly increased infarct volume and impaired cerebral blood flow recovery 24h after ischemic stroke in females. 45-minute MCA occlusions were performed on EC Panx1 overexpressing and control mice, which were confirmed with reduced CBF, measured by LSCI, at the end of the MCA occlusion in both sexes. (**A and C).** Infarct volumes were quantified using TTC staining in females **(B and E)** and males **(D and F).** CBF was measured using LSCI 24 hr prior to MCAO (baseline), at the end of the MCA occlusion (MCAO), and 24 hr post-MCAO/reperfusion in females **(G)** and males **(H)**. EC Panx1 Intact: N = 8 females and 8 males. EC Panx1 OvExp: N = 7 females and 9 males. Student’s T test (**A-D**) and Two-way ANOVA/Sidak’s multiple comparisons post hoc test (**G and H**). *p<0.05; **p<0.01; ****p<0.0001.

## Discussion

In the current study, we examined the effects of overexpression of EC Panx1 to evaluate if EC Panx1 content dictates cerebral arterial function and subsequent ischemic stroke recovery. Use of the ROSA26 locus to express the human isoform of Panx1^18^ in our Cdh5-Cre^ERT2+^ mice resulted in significantly increased Panx1 expression in cerebral endothelial cells in both male and female mice (**Fig 1**). Overexpression of EC Panx1 did not alter peripheral vascular function, including myogenic reactivity in mesenteric arteries or systemic blood pressure and heart rate in both male and female mice (**Fig 2**). However, overexpression of EC Panx1 increased posterior cerebral artery myogenic tone development due to reduced active diameter with no change in passive diameter only in female mice (**Fig 3A, C, and E**). Our data do suggest a small but significant reduction in passive diameter of the posterior cerebral arteries of male mice (**Fig 3F**), although this change does not result in a change in myogenic reactivity (**Fig 3B**) or a change in recovery to injury as male mice showed no change in cerebral blood flow recovery or infarct volume 24 hr post-ischemia/reperfusion injury (**Fig 4D, and F**). Overexpression of EC Panx1 only altered ischemic stroke outcome in female mice (**Fig 4**). Infarct volume 24 hr post-ischemia/reperfusion injury was significantly increased and recovery of cerebral blood flow in the ischemic hemisphere was significantly reduced in female mice (**Fig 4**). Together these data suggest that overexpression of EC Panx1 specifically in females results in increased vascular tone of cerebral arteries, which prevents sufficient recovery of cerebral blood flow into the ischemic region and results in a larger infarct area.

Although purinergic signaling and Panx1 channels had previously been identified as major regulators of ischemic stroke outcome^19–21^, we were the first to examine the role of vascular Panx1 in cerebral vascular physiology and ischemic stroke outcome^11^. Pharmacological inhibition of Panx1 with non-specific small molecule inhibitors has been shown to reduce ischemic stroke infarct volume^22–27^. We previously demonstrated that pharmacological inhibition of Panx1 with multiple small molecule inhibitors or a mimetic peptide significantly reduced cerebral artery myogenic tone development^11^. Therefore, our initial study examined the impact of SMC verse EC deletion of Panx1 on cerebral vascular function as Panx1 is expressed in both SMC and EC of cerebral arteries^11, 15–17^. Our results demonstrated that deletion of SMC using the SMMHC-Cre Panx1 floxed mice, did not alter cerebral myogenic reactivity or ischemic stroke outcome^11^. However, deletion of EC Panx1 reduced cerebral myogenic tone development and reduced infarct volume 24 hr post-MCAO/reperfusion^11^. Our studies here focused on the impact of overexpression of EC Panx1; however, it remains unknown if overexpression of SMC Panx1 has an impact on cerebral vascular function and ischemic stroke outcome.

Purinergic signaling can regulate both vasoconstriction and vasodilation in arteries^13, 14^. Activation P2X and P2Y receptors, which are activated by extracellular purine nucleotides, have been found to play a role in cerebral artery vasoconstriction^10, 28^. Knockdown or inhibition of P2Y4 and P2Y6 results in impaired myogenic tone development in cerebral arterioles^10^. Activation of P2X1 and P2X4 can also cause vasoconstriction in rat middle cerebral arteries^28^. Hydrolyzation of extracellular purine nucleotides by ectonucleotidases results in nucleotide formation such as adenosine, which can then promote vasodilation through adenosine receptors^13^. Deletion of CD39, an ectronucleotidase, results in impaired cerebral blood flow in the ischemic hemisphere 24hr post-ischemic stroke^29^. Taken together, these studies suggest an isoform-specific role of purinergic receptors in determining cerebral vascular tone. Our previous data indicated that SMC Panx1 was not involved in regulating myogenic tone in healthy cerebral arteries, but EC Panx1 content dictates extent of myogenic reactivity^11^. However, it remains unclear how EC Panx1 is regulating purinergic signaling within the vasculature to alter cerebral vascular tone or if Panx1’s role is independent of purinergic receptors. One hypothesis is that EC Panx1-mediated ATP release in response to increased luminal pressure activates SMC purinergic receptors resulting in pressure-induced vasoconstriction. Further studies are necessary to elucidate the downstream mechanism by which EC Panx1 contributes to myogenic tone development in cerebral arteries.

Our data demonstrate a sex-specific effect of overexpression of EC Panx1 where only females demonstrated impaired cerebral blood flow recovery 24h post-ischemic stroke and larger infarct volumes compared to control females (**Fig 4B and E**), which may be due to the increase in myogenic tone observed in female cerebral arteries at baseline (**Fig 3A**). Only two studies in the literature included females in examining the role of Panx1 in ischemic stroke outcome^30, 31^, and only one of those studies was appropriately powered to examine sex specific differences^30^. This study found that Panx1 expression in the brain is increased in female mice compared to male mice 4 days following a permanent middle cerebral artery occlusion, however the increase was minimal^30^. Additionally, similar to our data from isolated EC (**Fig 1D**), expression of Panx1 in healthy whole brain was not found to be different between males and females^30^. Freitas-Andrade et al also found that global knockout of Panx1 or inhibition of Panx1 with probenecid was only beneficial to females, resulting in reduced infarct volume and reduced number of Iba1+ and GFAP+ cells in female mice^30^. These data suggest that Panx1 may be a viable target to improve ischemic stroke outcome specifically in women.

Risk of ischemic stroke increases with age in both men and women, although women have an increased lifetime risk of stroke and increased stroke-related mortality^2^. Further, among stroke survivors, outcomes in women, including mobility limitation and anxiety or depression, are worse compared to men^1, 2, 32–35^. While some of these disparities may be due to social determinants, there are substantial differences in cardiovascular risk between men and women that change after menopause, suggesting a biologic contribution that may involve female sex hormones^2^. Estrogen has been shown to regulate cerebral artery myogenic tone development^36, 37^. Myogenic reactivity in cerebral arterioles is increased in mice with bilateral ovariectomy (OVX) and treatment with 17β-estradiol returns myogenic tone to ovary-intact female levels^36, 38^. Ovarian estrogen has been shown to be protective in ischemic stroke as demonstrated by OVX increasing infarct volume and estrogen replacement reducing infarct volume back to ovary-intact mouse levels^39, 40^. Multiple mechanisms have been suggested for the beneficial effects of estrogen post-ischemic stroke, including inflammatory processes, neurogenesis, and CBF recovery^41, 42^. Future studies are necessary to determine if EC Panx1 is regulated by or regulates estrogen-dependent signaling in the cerebral arteries.

Our findings suggest that increased expression of EC Panx1 may play a detrimental role in females following an ischemic stroke. Understanding disease states that may elevate Panx1 expression prior to the onset of a stroke is important for understanding which patients may benefit from early treatment with a Panx1 inhibitor post-ischemic stroke. Panx1 protein levels have been found to be increased in adipose tissue from humans with obesity and in skeletal muscle form obese mice compared to control mice^43, 44^. Panx1 mRNA expression has also been seen to increase in mouse models of abdominal aortic aneurysm (AAA), in lung tissue post-ischemia/reperfusion (IR), in kidney tissue post-IR, and in high shear stress areas of atherosclerotic lesions^45–47^. In isolated neurons, oxygen and glucose deprivation was shown to increase mRNA Panx1 expression, which was maintained with return of normal oxygen and glucose levels^48^. In addition to expression of Panx1, the open probability of the channel also plays an important role in regulating the physiological and pathophysiological consequences of Panx1 activation^49–51^. Phosphorylation of Panx1 in neurons in response to excitotoxic NMDA receptor signaling, which occurs during an ischemic injury, results in increased Panx1 channel activity without changes in overall expression^50^. Expression changes within individual cell types, including vascular SMC or EC, have not been examined with age, disease or injury. A limitation of our study is the use of young 4–6-month-old mice that are otherwise healthy. Future studies examining the expression changes of Panx1 within vascular cells during cardiovascular associated diseases and the impact of Panx1 on ischemic stroke outcome with co-morbidities are necessary as cardiovascular diseases increase the risk of ischemic stroke^1^.

Long-term CBF changes have been observed in patients following an ischemic stroke and hypoperfusion is associated with worse neurological function post-ischemic stroke^9^. Cerebral arteries are estimated to contribute ∼50% of cerebral vascular resistance and are largely regulated by myogenic reactivity, where increases in luminal pressure result in vasoconstriction, which is vital for autoregulation and control of CBF^7, 8^. Following a cerebral ischemia/reperfusion injury, myogenic tone is increased in cerebral arterioles^52^, which may reduce reperfusion into the ischemic region and impact infarct size. Given the limited therapeutic intervention for ischemic stroke and projected increase in global incidence of ischemic stroke, examination of the mechanisms contributing to altered myogenic reactivity post-ischemic stroke is important for identification of novel ischemic stroke therapeutic targets. Our previous^11^ and current data support a critical role of endothelial cell (EC) Panx1 in dictating cerebral artery function and ischemic stroke outcome. Furthermore, the existence of multiple FDA approved small molecule inhibitors of Panx1, including spironolactone^53^ and probenecid^54^, suggests that EC Panx1 may provide a novel pharmacological therapeutic target to improve post-ischemic stroke outcomes, in particular for women, a critically unmet medical need.

## Methods

### Sex as a biological variable

Both sexes of mice were used throughout the study and analyzed and compared individually.

### Mice

Male and female mice were used between 14-24 weeks of age and were fed normal chow and housed under a 12-hour light/dark cycle. Cdh5-Cre^ERT2+^ mice were crossed with our ROSA26-human Panx1 (hPanx1) transgenic mice^18^ to create a mouse that conditionally overexpresses human Panx1 (hPanx1^Tg^) in endothelial cells (EC). The ROSA26-hPanx1*^fl-STOP-fl^* transgene contains a floxed stop codon followed by the human Pannexin1, which has a FLAG-tag to allow identification of the transgene-encoded human Panx1 protein. To induce the overexpression of the hPanx1^Tg^ gene, 6- to 8-week-old mice received daily injections of tamoxifen for 10 days (1 mg/kg/day diluted in peanut oil; Sigma^11^). This deletes the stop cassette and results in expression of hPanx1^Tg^ in EC. Cre^−^ R26-Panx1^Tg^ were used as controls, which also received tamoxifen injections.

### Immunochemistry

Posterior cerebral arteries (PCAs) were dissected, perfused, PFA fixed, and paraffin embedded as previously described^11^. Briefly, 5 µm cross sections were subjected to paraffin removal and washing, antigen retrieval (antigen unmasking solution, Vector), blocking for 30 minutes and stained with Flag tag (Cell Signaling; DYKDDDDK Tag (D6W5B); 1:100). Sections were mounted in DAPI containing ProLong Gold Antifade Mountant (Invitrogen) and imaged using an Olympus FluoView 1000.

### EC isolation for cerebral EC protein and western blot analysis

Protein was from harvested EC isolated from the brain. EC were isolated from brains using the Adult Brain Dissociation Kit, mouse and rat (Miltenyi), following a modified manufacturers protocol. Briefly, brains were collected and minced into the Enzyme mix 1 (Buffer Z and Enzyme P), rotated at 37°C for 17 min at 20 RPM, then Enzyme mix 2 (Buffer Y and Enzyme A) was added and the sample mechanically dissociated using a Pasteur pipette. The solution was then rotated at 37°C for 12 min at 10 RPM, mechanically dissociated by passing through a 20 g needle 10 times, and finally rotated at 37°C for an additional 10 min at 10 RPM. The dissociated brain tissue was passed through a 70 µm filter followed by 10 mL HBSS with divalent cations and MACS Buffer. Myelin was then removed using a 25% density gradient solution (Percoll; Sigma) and RBCs were removed using the RBC lysis solution (Miltenyi). Endothelial cells were isolated using CD31 microbeads (1:10), LS columns, and the MACS Separator (Miltenyi) into MACS Buffer. After centrifugation, cerebral endothelial cell protein was isolated in RIPA lysis buffer. Samples were analyzed for protein expression using western blotting by running on samples on SDS-Page gels, transferred to nitrocellulose membranes and then stained with Panx1 at (Abcam; 1:1,000) and Beta-Tubulin (Cell Signaling; 1:1,000). LICOR Odyssey M was used for scanning of membranes and protein quantification.

### Pressure myography

Myogenic tone development was measured in PCA and third order mesenteric arteries using pressure myography as previously described^11^. Cerebral vessels were isolated from euthanized mice, cannulated on glass micro-pipets, placed in a pressure myography chamber (Living Systems and Danish Myo Technology) containing Krebs-HEPES solution (in mM; 118.4 NaCl, 4.7 KCl, 1.2 MgSO_4_, 4 NaHCO_3_, 1.2 KH_2_PO_4_, 2 CaCl_2_, 10 Hepes, and 6 glucose), and equilibrated to 37°C. The lumen was filled with the same Krebs-HEPES solution and maintained in a no-flow state to equilibrate for 30 minutes at 40 mmHg of intraluminal pressure. Myogenic reactivity was measured by reducing the intraluminal pressure to 20mmHg and incrementally increasing it in 20mmHg steps to a maximal pressure of 100 mmHg (PCA) or 120 mmHg (mesenteric arteries). At each step, the active lumen diameter is recorded after a 5-minute stabilization period. EC health was confirmed by >70% dilation to 5 µM NS309 and SMC health was confirmed by >30% constriction to 30 mM KCl. The Krebs-HEPES solution was then replaced with a Ca^2+^ free Krebs-HEPES supplemented with 2 mM Ethylene glycol-bis(2-aminoethylether)-N,N,N’,N’-tetraacetic acid (EGTA; Sigma) and 10 μM sodium nitroprusside (Sigma), and the vessel was equilibrated at 80 mmHg intraluminal pressure for 30 minutes. The passive lumen diameters were measured during incremental increases of lumen pressure. Myogenic tone was calculated at each pressure step using the following equation: % myogenic tone = [(passive diameter – active diameter)/passive diameter] x 100.

### Laser speckle contrast imaging (LSCI) for cerebral blood flow (CBF) measurement

LSCI (Perimed) was used to measure CBF at baseline, which was performed 24 hr before surgery, at the end of the 45 min occlusion while the filament is still in place, and 24 hr post-reperfusion. To measure CBF, fur was removed from the scalp using Nair (only necessary for baseline measurement) and then an incision was made in the scalp to reveal the skull. Mice were maintained under isoflurane anesthesia set as low as possible (1.0-1.5%) and blood flow was recorded within 5 minutes. After recording, the scalp skin was sutured, and mice were removed from anesthesia and placed back to their cage with access to water and food. Bupivacaine was administered prior to incision for topical analgesic relief.

### Middle Cerebral Artery (MCA) Occlusion (MCAO)/Reperfusion

Ischemic stroke injury was achieved performing an MCAO with a monofilament as previously described^11^. Mice were anesthetized with 1.5-2% isoflurane and a midline incision was made at the ventral surface of the neck revealing the left external carotid artery. A 6-0 medium MCAO suture L34 reusable monofilament with a round tip (Doccol Crop.) was inserted through the external carotid artery into the internal carotid artery and pushed up to the MCA until resistance is felt and occlusion achieved. The neck wound is sutured closed and the mouse is then removed from anesthesia. After 45 minutes, the occlusion was confirmed using LSCI to measure CBF, where ischemia was defined as a > 30% and < 60% reduction in CBF of the ischemic hemisphere compared to baseline CBF. The monofilament was then removed to allow for reperfusion, and the mouse was placed back into its cage with access to water and soft food. Body temperature was maintained at 37°C during surgery and for 2 hr post-reperfusion. Bupivacaine was administered prior to incision in the necks for topical analgesic relief and buprenorphine was administered 1 hr post-reperfusion.

Infarct volume was analyzed 24 hours after reperfusion. Mice were euthanized by cardiac exsanguination and brains were harvested. 2mm sections were stained with 2,3,5-Triphenyltetrazolium chloride (TTC) (Sigma; 2%) to visualize the ischemic region. Infarct volume was quantified using ImageJ, correcting for edema, with the following calculation: % infarct volume = [(non-ischemic hemisphere area – (ischemic hemisphere area – infarct area)) * 100]/ non-ischemic hemisphere area. Successful ischemic injury was defined as > 5% and < 60% infarct volume.

### Blood pressure through radiotelemetry

12-week-old mice were implanted with radiotelemetry devices from Data Sciences International (DSI) as previously described^11^. Under isoflurane anesthesia, the catheter of the radiotelemetry unit was placed in the left carotid artery and advanced towards the heart until resistance was felt. The radiotransmitter was then placed in a subcutaneous pouch on the right flank. Mice were allowed to recover for 7 days and then blood pressure was continuously recorded for 5 days. **Statistics**. Data were analyzed using Prism 10 software (GraphPad Software Inc.) and displayed in mean ± SEM. Students t-tests, one-way or two-way ANOVA (with repeated measures) were used as necessary and specified within figure legends. * p < 0.05, ** p < 0.005, *** p < 0.0005

### Study approval

All studies were conducted in accordance with the Tufts Medical Center IACUC.

### Data availability

Data is available upon request to the corresponding author.

## Author Contributions

MTG conducted experiments, acquired, and analyzed data, and wrote the manuscript. AKM, CKD, EYD, and GS conducted experiments and analyzed data. CM and KR created and provided the ROSA26-hPanx1 Tg mice. BEI contributed to the design and execution of the research studies. MEG designed the research studies, conducted experiments, analyzed data, and wrote the manuscript. All authors contributed to reviewing and editing the manuscript.

## Acknowledgements

This work was supported by NIH HL143165 (MEG) and NIH P01 HL120840 (BEI and KSR). We would like to thank the MCRI Small Animal Physiology Core Lab, which provided space and support for the included experiments.

